# Characterization and application of hyperthermia-evoked seizures in a mouse model of focal cortical dysplasia

**DOI:** 10.64898/2025.12.19.695581

**Authors:** Xiaogang Zhang, Chandni Rana, Ruitao Shen, Tao Yang, Joseph Barden, Hudson Haberland, Mi Jiang, Rida Qureshi, Saif Siddiqui, Alexander Sajdak, Patrick Lawlor, Yu Wang, Joanna Mattis

## Abstract

Rodent models of drug-resistant epilepsy are widely used to uncover mechanisms of seizures and to test the efficacy of treatments; however, the stochastic and relatively infrequent nature of spontaneous seizures in these models makes it challenging to observe and manipulate peri-ictal activity, and to rapidly screen therapeutics. As fever has been identified as a clinical seizure trigger, we investigated the ability of induced hyperthermia to trigger seizures in a mouse model of focal cortical dysplasia (FCD), a common cause of drug-resistant epilepsy. Experimental mice were generated via *in utero* electroporation of a gain-of-function RHEB variant (RHEBp.P37L). In adult *Rheb-*FCD mice, hyperthermia elicited behavioral seizures ∼80% of the time, with a mean temperature threshold of ∼40 °C. Using this acute paradigm, as a proof-of-concept, we found that chemogenetic inhibition of a subset of excitatory neurons within the dysplasia significantly elevated the threshold for hyperthermia-evoked seizures. We then confirmed the relevance of this acute finding to spontaneous epileptiform activity, using repeated chemogenetic inhibition across chronic recordings to confirm a reduction of spontaneous seizures as well as interictal spikes. Overall, our data demonstrate that hyperthermia-evoked seizures can be readily elicited in *Rheb-*FCD mice, offering a tractable platform for evaluating therapeutic interventions targeting seizure susceptibility in focal epilepsy.

## Introduction

Epilepsy is a highly prevalent neurological condition that imparts a significant worldwide burden as measured by disability-adjusted life years (Fiest et al., 2017). Approximately one-third of persons with epilepsy have drug-resistant disease, defined as continued seizures despite adequate trials of two appropriately selected and tolerated antiseizure medications (ASMs) (Kwan et al., 2010; Ramos-Lizana et al., 2012; Chen et al., 2018). There is therefore an urgent need to improve our mechanistic understanding of refractory epilepsies and to develop new therapeutic interventions with improved efficacy.

Rodent models are an indispensable experimental platform for dissecting the mechanisms underlying epilepsy and for conducting preclinical ASM screening (Marshall et al., 2021). These models generally fall into two broad categories, each with distinct methodological and translational advantages. First, acute seizures can be evoked in wildtype (WT) animals by a range of methods, including electroshock, kindling, or chemoconvulsants (e.g. (Leech and McIntyre, 1976; Schmidt, 1987; Karler et al., 1989; Cavalheiro, 1995; Giardina and Gasior, 2009; Lévesque and Avoli, 2013)). These seizures provide controlled, acute assays for assessing seizure thresholds and evaluating ASM efficacy but may be mechanistically dissimilar to spontaneous seizures as occur in epilepsy. Second, spontaneous seizures can be recorded across a range of models of chronic epilepsy, including sodium channelopathies (Griffin et al., 2018; Meisler, 2019; Scott et al., 2025), absence epilepsy (Jafarian et al., 2020), temporal lobe epilepsy (TLE) (Lévesque et al., 2016), and focal cortical dysplasia (FCD) (Nguyen and Bordey, 2022). These chronic epilepsy models have greater construct validity, but the unpredictability and stochasticity of spontaneous seizures pose significant challenges for experiments requiring acute observation and manipulation, and necessitate a low-throughput, time-intensive approach for the screening of potential new therapeutics.

Body temperature elevation (i.e., fever) is a well-established seizure trigger in persons with epilepsy. This phenomenon is most notably observed in Dravet Syndrome (DS), a severe developmental and epileptic encephalopathy in which febrile seizures constitute a cardinal diagnostic criterion (Dravet and Oguni, 2013; Connolly, 2016). Consistent with this, experimentally-induced hyperthermia is commonly employed to elicit seizures in mouse models of DS (e.g., (Oakley et al., 2009; Mistry et al., 2014; Mattis et al., 2022)). Even beyond DS, fever – whether in the context of systemic illness or other immune response – ranks among the most frequently recognized triggers of breakthrough seizures in persons with epilepsy (Bauer et al., 2000; Balamurugan et al., 2013; Kaddumukasa et al., 2013; Illingworth et al., 2014; Wassenaar et al., 2014; Kurian et al., 2018; Rafati et al., 2023) and hyperthermia has also been occasionally employed as a seizure trigger in rodent models of PCDH19-related epilepsy (Cwetsch et al., 2022) and Angelman syndrome (Gu et al., 2019).

Clinical case reports indicate that fever can trigger seizures in patients with FCD (Singh et al., 2001; Itamura et al., 2019), a common cause of drug-resistant focal epilepsy (Harvey et al., 2008; Crino, 2009; Guerrini and Dobyns, 2014; Blumcke et al., 2017). We reasoned that if hyperthermia could be used to evoke seizures in a mouse model of FCD – a model which otherwise recapitulates key pathophysiological and electroclinical features of the clinical condition –– this would provide an acute focal-onset seizure model that was experimentally tractable and that offered high construct, face, and predictive validity.

In this study, we first established hyperthermia as a reliable and reproducible method for eliciting seizures in a RHEBp.P37L-based model of focal cortical dysplasia type II (*Rheb*-FCD) (Yan et al., 2006; Hsieh et al., 2016; Nguyen et al., 2019). As a proof-of-concept, we then utilized this experimental platform to confirm that inhibition of excitatory neurons within the dysplastic region provided protection against both hyperthermia-evoked and spontaneous seizures. Our findings introduce a novel methodological framework for triggering acute seizures in a focal epilepsy model and demonstrate excitatory neurons as a critical cellular target for potential intervention in FCD.

## Methods

### Experimental mice

To generate the *Rheb-*FCD mouse model, *in utero* electroporation (IUE) was performed at embryonic day (E)14.5-15 to express RHEBp.P37L, a patient-specific gain-of-function pathogenic mutation (Proietti Onori et al., 2021), in the forebrain neural progenitors. pCAG-RhebP37L (1.0 µg/µL) was co-injected with GFP (1.0 µg/µL) or CaMKIIα-hM4DGi-mCherry / pCAG-GFP (1.0 µg/µL and 0.5 µg/µL, respectively) vectors. When applicable, control animals were generated (in the same pregnant dam) using a red fluorescent plasmid so that they could be easily distinguished from *Rheb-*FCD mice by fluorescent illumination shortly after birth. Electroporation was performed on mice of either a CD1 background (RRID: IMSR_CRL:022) or C57BL/6 background (RRID: IMSR_JAX:000664), as specified.

After induction of anesthesia with isoflurane (4-5% induction and 2-4% for maintenance), the pregnant dam was subjected to an abdominal incision to enable visualization of the embryos. Plasmids were injected into the lateral ventricle and electroporated into the neural progenitor cells along the dorsal telencephalon via a pulled glass micropipette. The electroporation was conducted across the embryonic head using five electric pulses (35 V, 50 ms duration, 1-second intervals, Boston apparatus, BTX-100). During the entire procedure, exposed embryos were kept moisturized with warm saline. After electroporation or injection, the uterus was placed back in the abdominal cavity, and the peritoneal cavity of the dam was lavaged with 5 mL of 0.9% saline containing an antibiotic-antimycotic. The surgical incision was then sutured closed.

To generate *Scn1a^+/−^* mice for experimental use, male mice on a 129S6/SvEvTac (129S) background that were hemizygous for the *Scn1a^tm1Kea^*/Mmjax knockout allele (129S.Scn1a^+/−^, [RRID:MMRRC_037107-JAX](https://www.mmrrc.org/catalog/sds.php?mmrrc_id=37107)) were crossed with female C57BL/6J (B6) mice. The hemizygous *Scn1a* progeny from this cross (i.e., *Scn1a^+/−^* mice on a 50:50 129S:B6 background) have been established to exhibit a well-characterized epileptic phenotype with spontaneous seizures as well as seizures that can be evoked by hyperthermia (Mistry et al., 2014).

### Hyperthermic seizure generation

Hyperthermia-evoked seizures were elicited as described previously (Mattis et al., 2022; Kravchenko et al., 2023). Mice were heated gradually using a heat lamp placed over the recording chamber. Internal body temperature was monitored continuously via a rectal probe connected to a BAT-12 thermometer box (Physitemp). Heating continued either until a behavioral seizure was observed or until internal body temperature reached 42.5 °C, at which point the lamp was removed and the mouse was cooled using ice.

To further increase experimental throughput, and to minimize handling of *Rheb-*FCD mice (which we anecdotally observed to trigger seizures in a subset of mice), for some experiments we placed the rectal probe in a control mouse – age, sex, and genotype-matched to the experimental mouse – which was heated simultaneously with the experimental mouse within the same chamber; in these cases we then used the internal body temperature of the “probe mouse” as a proxy for that of the *Rheb-*FCD experimental mouse.

### Intracranial electroencephalography (EEG) recording and analysis

Mice were anesthetized by isoflurane inhalation and placed in a stereotaxic apparatus (Kopf Instruments). Craniotomies were made using a high-speed stereotaxic drill (Kopf Instruments). Three stainless steel screw electrodes (#8IE3639616XE; P1 Technologies) were implanted with 2 electrodes over the left and right parietal lobes and a reference electrode over the cerebellum. The electrode sockets were connected to a 6-pin pedestal (#8K000229801F; P1 Technologies) and secured to the skull with dental cement. Mice were allowed to recover for at least 3-5 days prior to monitoring.

EEG signal was collected using Natus NeuroWorks (Natus Medical Inc.) software and acquisition hardware. Signals were sampled at 4096 Hz. Custom Python code (https://github.com/mattis-laboratory/fcd-hyperthermia) was developed for the detection of interictal spikes and seizures.

Interictal spike detection was performed to quantify epileptiform activity between seizures. Data were filtered with a 60 Hz notch filter and a 1-70 Hz second-order Butterworth bandpass filter. To establish a stable baseline for normalization, an initial 30-minute window was selected from the baseline period of each experiment. This window was subdivided into six consecutive 5-minute segments. The standard deviation of each segment was calculated, and the coefficient of variation (CV) across the standard deviation of the segments was computed as *CV* = (σ_*segments* / μ_*segments*) × 100. If CV exceeded 20%, indicating instability in the baseline, the entire 30-minute window was iteratively advanced by 5 minutes and re-evaluated until CV ≤ 20% was met or the search reached the experimental timeperiod. The entire experimental epoch was then z-scored using the mean and standard deviation of this validated baseline period. Spike detection was performed on the z-scored signal using scipy’s find_peaks function. Because some interictal spikes manifested as positive deflections, and others as negative deflections, we performed spike detection separately for positive and negative deflections and then combined the results in a manner that avoided duplicate counting (see below). The spike detection criteria included: (1) amplitude ≥ 4.5 standard deviations above baseline mean, (2) width 20-200 ms, (3) prominence ≥ 2.25 standard deviations, and (4) minimum interspike interval (ISI) of 20 ms. Detected spikes were filtered to exclude (1) artifacts within 5% of the recording edges, (2) excessively large artifacts, (3) spikes occurring during previously identified seizure periods (see below). Duplicate detections within the minimum ISI were resolved by retaining the event with higher prominence.

Electrographic seizures were defined on EEG as repetitive epileptiform discharges at > 2 cycles/second and/or a characteristic pattern with spatiotemporal evolution (i.e., gradual change in frequency and amplitude) lasting at least 10 seconds (Kao et al., 2023). Spontaneous seizures were identified by an initial manual review of the entire EEG, confirmed by a second, focused review of EEG epochs flagged by a custom-coded automated seizure detector. This detector, developed in Python using MNE-Python (an open-source Pyhton package for visualization and analysis of neurophysiological data) and SciPy, was designed to identify sustained periods of high-amplitude activity. For each selected channel, data were down sampled (from 4096 Hz to 512 Hz) to lower data size while preserving signal integrity, filtered (60 Hz notch filter;1-70 Hz fourth-order Butterworth bandpass filter), and z-scored. The 97^th^ percentile of the absolute z-values was used as a threshold for detecting unusually strong activity. The data were analyzed in one-second windows, and the proportion of samples above the threshold within each window was calculated. Windows were marked as “active” when at least 10% of samples exceeded the threshold. Consecutive active windows lasting at least seven seconds were considered candidate seizure epochs, and events separated by gaps of 30 seconds or less were merged to avoid splitting longer events. Each detected epoch was then plotted to facilitate manual review for confirmation.

### Video recording and analysis

Video recordings were manually reviewed to quantify the duration, severity, and characteristics of evoked seizures. Seizure onset was defined as the first instance of sustained, rhythmic, and abnormal motor activity that could not be reasonably attributed to normal baseline, exploratory, or grooming behavior. The severity of each event was rated using a modified Racine scale (Erum et al., 2019). We restricted our analysis to behaviors corresponding to Racine stages 3 through 7, as these were generally unambiguous. All scoring was completed by two trained observers blinded to the experimental condition.

### Chemogenetics

For chemogenetic inhibition experiments, clozapine N-oxide (CNO; CAS Number: 34233-69-7) (5mg kg^−1^) was diluted in 2% dimethyl sulfoxide (DMSO) and injected intraperitoneally. An equivalent volume of vehicle solution (2% DMSO in sterile saline) was injected as a control. For acute experiments, injections were performed 30 minutes prior to hyperthermia. For chronic experiments, a single injection was performed once daily (at 10am) for 6 consecutive days, and EEG was analyzed within a window of 30 minutes to 6 hours post injection. All experiments utilized a cross-over design: on the first day, each mouse was randomly assigned to receive either CNO or vehicle; on the second day, mice that had received CNO received vehicle, and vice versa.

### Immunohistochemistry

For c-Fos immunostaining, mice were deeply anesthetized with isoflurane and transcardially perfused with 5 mL of ice-cold PBS followed by 5 mL of 4% paraformaldehyde (PFA). Brains were removed and postfixed at 4 °C overnight in 4% PFA, then equilibrated overnight in 30% sucrose in PBS. Tissue was sectioned at a thickness of 40 μm using a frozen microtome (Leica Biosystems, SM2010) and sections were placed in cryoprotectant (25% glycerol, 30% ethylene glycol in PBS, pH adjusted to 6.7 with HCl) for storage at −20 °C. Immunofluorescence staining was performed on free-floating sections, which were washed with PBS (3 washes, 5 minutes each), blocked and permeabilized for 1 hour at room temperature in PBS containing 0.3% Triton X-100 and 3% normal donkey serum (NDS), and then incubated overnight at 4 °C with rabbit anti-c-Fos (Cell Signaling Technology, RRID: AB_2247211) diluted 1:500 in the same solution. Following primary antibody incubation, sections were again washed with PBS (3 washes, 5 minutes each) and incubated for 3 hours at room temperature in donkey anti-rabbit Alexa Fluor 647 (Thermo Fisher Scientific, RRID: AB_2492288) diluted 1:500 in PBS containing 0.3% Triton X-100 and 3% NDS. Sections were then washed in PBS, incubated in 1:50,000 4,6-diamidino-2-phenylindole (DAPI; Thermo Fisher Scientific, RRID: AB_2307445)) in PBS at room temperature for 15 min, and washed a final time in PBS. Sections were mounted and cover-slipped with PVA-DABCO mounting medium (MilliporeSigma, RRID:SCR_024103).

For pS6 immunostaining, the experimental procedure was similar except as follows: Mice were anesthetized via brief exposure to isoflurane followed by an overdose of pentobarbital. Tissue was sectioned at a thickness of 70 μm using a frozen microtome (Leica Biosystems, VT1000S). Blocking and antibody incubations were in PBS containing 0.2% Triton X-100 and 5% NDS. Primary antibody was rabbit anti-pS6 (Cell Signaling Technology, RRID:AB_331679) diluted 1:500 and incubated overnight at room temperature. Secondary antibody was donkey anti-rabbit Alexa Fluor 594 (Invitrogen, RRID:AB_141637) diluted 1:500 and incubated for 1-2 hours at room temperature. Sections were mounted using Glycergel mounting medium (Agilent, #C056330-2), air-dried, and sealed with coverslips.

### Imaging and analysis

Fluorescence images were acquired on a Nikon ECLIPSE Ni microscope controlled by Nikon NIS-Elements software. Imaging parameters were kept constant across animals, sessions and experimental groups for each experiment.

c-Fos positive cell counts were performed using Fiji (ImageJ). The GFP (Rheb+) channel served as the anatomical reference for the dysplastic region. For each section, a region of interest (ROI) was manually delineated to encompass the entire dysplastic area while maintaining comparable areas across sections. Within the ROI, the far red (c-Fos) channel was processed by (1) smoothing to suppress high frequency noise (“smooth filter), (2) thresholding to produce binary masks that occupied ∼10% of pixels (“moments” algorithm in B&W mode), and (3) segmenting to separate adjacent nuclei (“watershed”). Cell counts were then obtained using “analyze particles,” using a minimum size of 30 µm^2^ and a circularity range from 0.00 to 1.00.

### Statistics and data display

Statistical analyses were performed using GraphPad Prism version 10.4.1 for Windows, GraphPad Software, Boston, Massachusetts USA, www.graphpad.com. GraphPad Prism was also used to generate all graphs. Schematics were prepared using BioRender software (www.BioRender.com).

## Results

### Seizures in *Rheb-*FCD mice are readily evoked by hyperthermia

To generate *Rheb-*FCD mice, we performed IUE to introduce a constitutively active, gain-of-function RHEB (RHEBp.P37L) variant known to induce mTOR hyperactivation (Proietti Onori et al., 2021) (Fig. 1A, Supplementary Fig. 1A-B). We confirmed that *Rheb*-FCD mice exhibit spontaneous seizures (Supplementary Fig. 1C-D), as previously shown (Yan et al., 2006; Hsieh et al., 2016; Nguyen et al., 2019).

**Figure 1.**
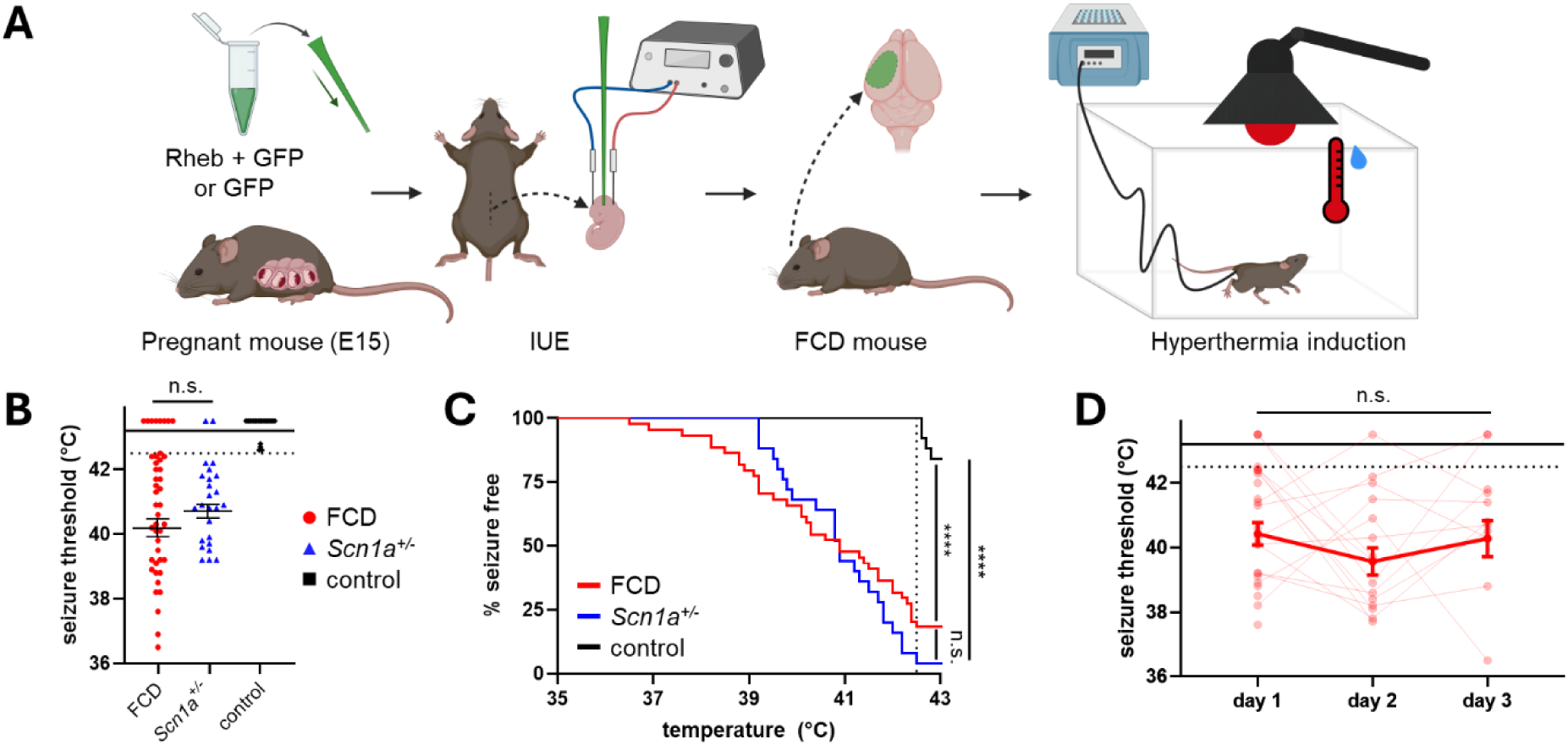
Seizures in adult *Rheb-*FCD mice are reliably evoked by hyperthermia. **(A)** Cartoon illustrating experimental overview for generating *Rheb-*FCD mice and eliciting seizures via hyperthermia. IUE-mediated somatic mutagenesis results in *Rheb-*FCD mice; control mice are generated via IUE to express a fluorophore. Hyperthermia was induced in adult mice via heat lamp exposure. **(B)** Hyperthermia-evoked seizure thresholds for *Rheb-*FCD mice (red circles), *Scn1a^+/−^* mice (blue triangles), and control mice (black squares). Symbols plotted above the horizontal solid line indicate sessions for which no behavioral seizure was observed. Seizures were elicited in 82% (36/44) of sessions for *Rheb-*FCD mice versus 96% (24/25) of sessions for *Scn1a^+/−^* mice and 16% (5/30) of sessions for control mice. Note that no control mice had seizures elicited prior to the experimental endpoint of active heating (42.5 °C; horizontal dashed line). For sessions that elicited seizures below the experimental endpoint, mean seizure threshold was similar between FCD and *Scn1a^+/−^* mice (40.2 ± 0.3 °C versus 40.8 ± 0.2 °C, p = 0.14, unpaired t-test). n = 24 mice / 44 sessions (FCD), 25 mice / 25 sessions (*Scn1a^+/−^*), 6 mice / 30 sessions (control). **(C)** Seizure thresholds across all sessions were plotted as a Kaplan-Meier curve to capture both threshold data and the percentage of mice that exhibited seizures. FCD and *Scn1a^+/−^* curves were similar (p = 0.31) and both were highly significantly different than control (p < 0.0001). Curves were compared using the log-rank (Mantel-Cox) Chi square test. **(D)** Seizure thresholds in *Rheb-*FCD mice across successive experimental days. Results from each individual mouse (light red) are indicated by lines connecting data points. Symbols plotted at the top of the graphs (i.e., above the horizontal solid line) indicate sessions for which no behavioral seizure was observed. Mean daily thresholds (dark red), calculated from trials that did evoke seizures, ranged from 39.6 ± 0.4 °C to 40.4 ± 0.4 °C and did not differ significantly across days (overall fixed effect p = 0.30, mixed effects model; pairwise comparisons 0.45-0.94, Tukey’s multiple comparisons test). n = 9-21 mice.

To determine whether hyperthermia could trigger acute seizures in adult (P35+) *Rheb*-FCD mice, we placed mice in a recording chamber and positioned a heat lamp above the chamber. The internal body temperature was continuously monitored, and animals were heated until either a behavioral seizure was observed, or until the experimental endpoint of 42.5 °C was reached. As we found no significant strain or sex differences in outcomes (Supplementary Fig. 2), we pooled data obtained across all *Rheb-*FCD mice. Littermates electroporated with a fluorophore aloe were included as a negative control. As a positive control, we included *Scn1a^+/−^*mice – in which the hyperthermic seizure phenotype is well established (Mistry et al., 2014) – for comparison. We found that seizures in *Rheb-*FCD mice were readily elicited by hyperthermia, with unambiguous behavioral seizures (Racine ≥ 3) observed in 82% (36/44) of sessions (n = 24 mice) at thresholds comparable to those observed in *Scn1a^+/−^* mice (40.2 ± 0.3 °C versus 40.8 ± 0.2 °C, p = 0.14, unpaired t-test) (Fig. 1B-C). Notably, no control mice exhibited seizures below the experimental endpoint of active heating (i.e., 42.5 °C). Seizure thresholds (among *Rheb-*FCD mice that exhibited seizures) were relatively stable across repeated seizure inductions performed on consecutive days (Fig. 1D).

To further characterize these hyperthermia-evoked seizures, we scored videos of the seizure sessions to determine the maximum Racine score (focusing only on the first hyperthermia session in CD1 mice; n = 19 mice). We observed that over half (11/19) of the cohort reached a maximum severity of Racine ≥ 5 (Fig. 2A). In a subset of mice (n = 8), we performed electrographic recordings to compare hyperthermia-evoked seizures with spontaneous seizures captured on continuous EEG. We found that the electrographic pattern (Fig. 2B) and duration (Fig. 2C) were similar for evoked versus spontaneous seizures.

**Figure 2.**
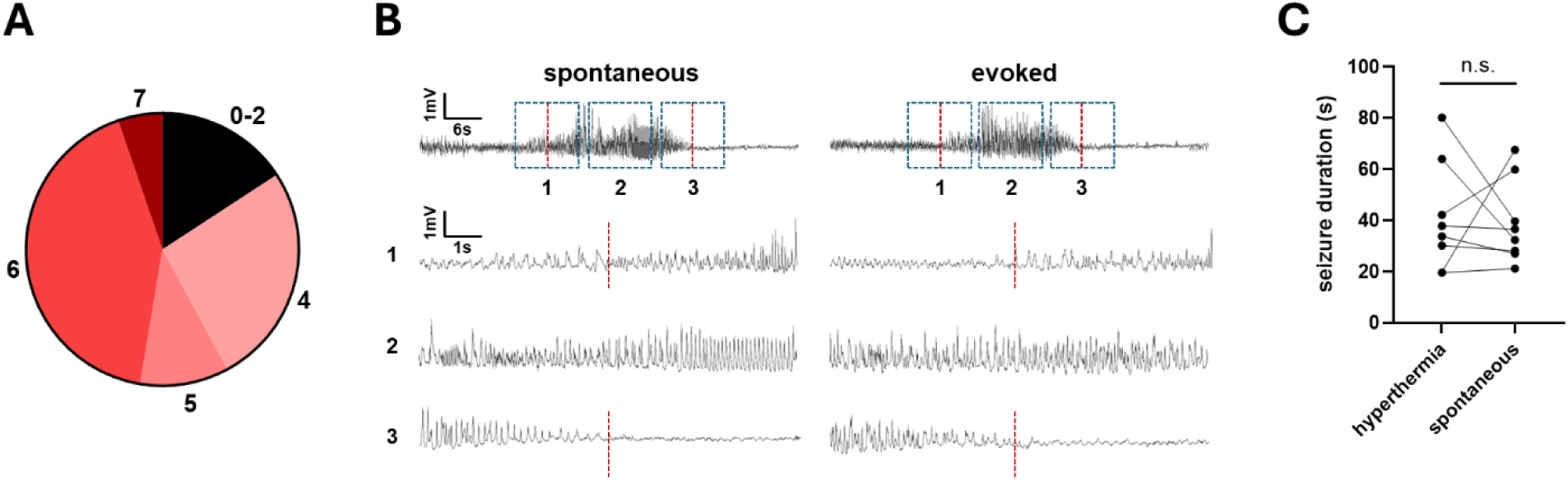
Electrographic and behavioral characterization of hyperthermia-evoked seizures in *Rheb-*FCD mice. **(A)** Maximum Racine score observed during hyperthermia-evoked seizures. A maximum seizure severity of Racine 4 was observed in ∼26% of mice, Racine 5 in ∼11%, Racine 6 in ∼42%, and Racine 7 in ∼5%. n = 19 mice. **(B)** Example EEG data recorded from the same animal during a spontaneous seizure (left) and a hyperthermia-evoked seizure (right). Vertical lines indicate seizure onset and offset. Regions spanning seizure onset (1), mid-seizure (2), and seizure offset (3) as indicated in the zoomed-out trace (top) are shown in the corresponding zoomed-in traces, below. **(C)** Electrographic seizure duration was similar between hyperthermia-evoked and spontaneous seizures (41 ± 7 seconds vs. 39 ± 6 seconds, p = 0.85, paired t-test). n = 8 mice.

Overall, these data demonstrate that hyperthermia can be used to reliably and repeatedly evoke seizures in adult *Rheb*-FCD mice, and that these seizures are behaviorally evident and are electrographically similar to spontaneous seizures. We therefore reasoned that the acute hyperthermia-evoked paradigm could be used to investigate mechanisms of seizures in the *Rheb-*FCD model.

### Chemogenetic inhibition of excitatory neurons within the dysplastic regions ameliorates both hyperthermia-evoked and spontaneous seizures

We next sought to demonstrate the experimental utility of the acute hyperthermia seizure induction method, employing it as a platform on which to test the efficacy of a potentially protective intervention. In patients with FCD, complete surgical resection of dysplastic tissue is a key positive prognostic factor for postoperative seizure freedom (e.g., (Cohen-Gadol et al., 2004; Chern et al., 2010; Jayalakshmi et al., 2019)), so we hypothesized that a spatially focused inhibition regionally targeted to the dysplasia would have a therapeutic, anti-seizure effect.

We began by confirming that hyperthermia-evoked seizures strongly activated neurons within the dysplasia, using expression of the immediate early gene, *Fos* (Hudson, 2018), as a readout. We used Fiji (ImageJ) to count the number of c-Fos positive cells within the dysplastic region, with GFP serving as the anatomical reference to define the ROI. We observed widespread c-Fos expression within the dysplasia that was significantly increased relative to anatomically-matched neocortex within control mice also subjected to hyperthermia (Fig. 3A-B). We also performed a within-mouse comparison of the FCD versus contralateral hemisphere among *Rheb-*FCD mice. Although contralateral c-Fos activation in *Rheb-*FCD mice was ∼38% higher than that observed in controls (mean c-Fos+ cells / ROI 4020 ± 290 versus 2922 ± 211) – as expected given that seizures may spread beyond the dysplastic hemisphere – we nevertheless observed significantly more c-Fos expression within the FCD relative to the contralateral hemisphere (Fig. 3C).

**Figure 3.**
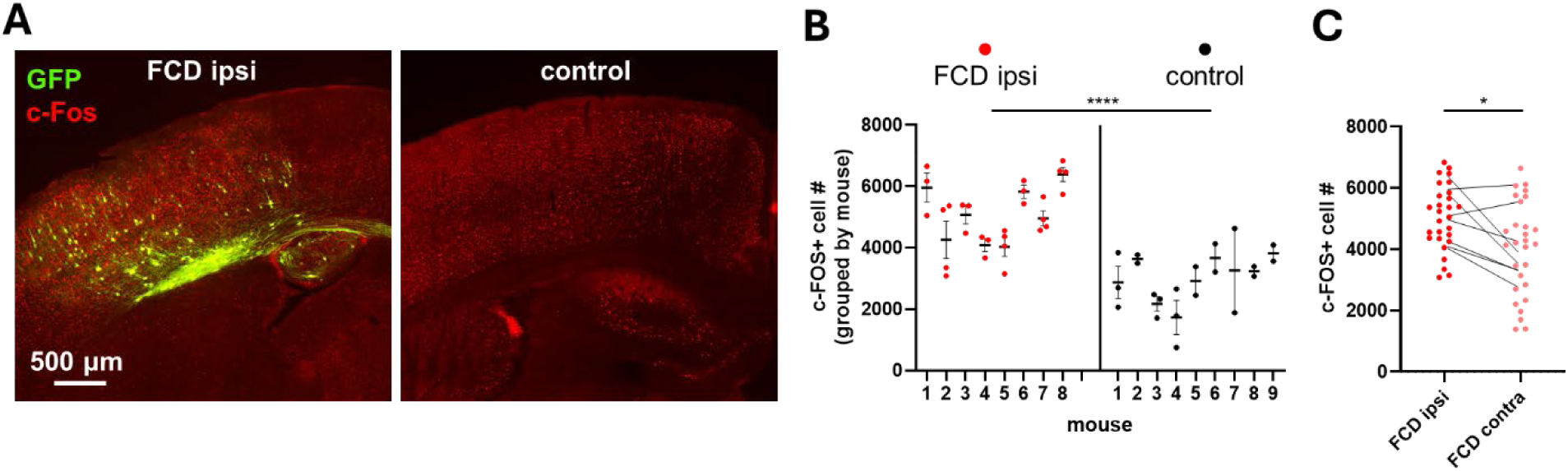
Hyperthermia-evoked seizures result in robust c-Fos activation within the dysplasia. **(A)** Representative images of seizure-associated c-Fos activation (red) within a dysplastic region (left; delineated by GFP expression (green)) versus within an anatomically matched neocortical region in a control mouse (right). **(B)** c-Fos+ cell number quantified ipsilateral to / within dysplastic regions (“FCD ipsi”) versus within anatomically matched neocortical regions in control mice. Each individual data point represents a count from one ROI; note that multiple ROIs were imaged per mouse. Nested unpaired t-test showed a significant difference between groups (p = 0.0001). **(C)** c-Fos+ cell number quantified within dysplastic regions (“FCD ipsi”) versus within the contralateral neocortex within the same mice (“FCD contra”). Each individual data point represents one ROI; lines connect mean counts (averaged across ROIs) for each individual mouse. Paired t-test showed a significant difference between regions (p = 0.03). n = 28 slices / 8 mice (FCD ipsi and contra); 21 slices / 9 mice (control).

We next sought to test whether chemogenetic inhibition of excitatory neurons within the region of dysplasia had a therapeutic effect in adult (P35-72) *Rheb-*FCD mice, taking advantage of the acute hyperthermia-evoked seizure experimental paradigm (Fig. 4A).

**Figure 4.**
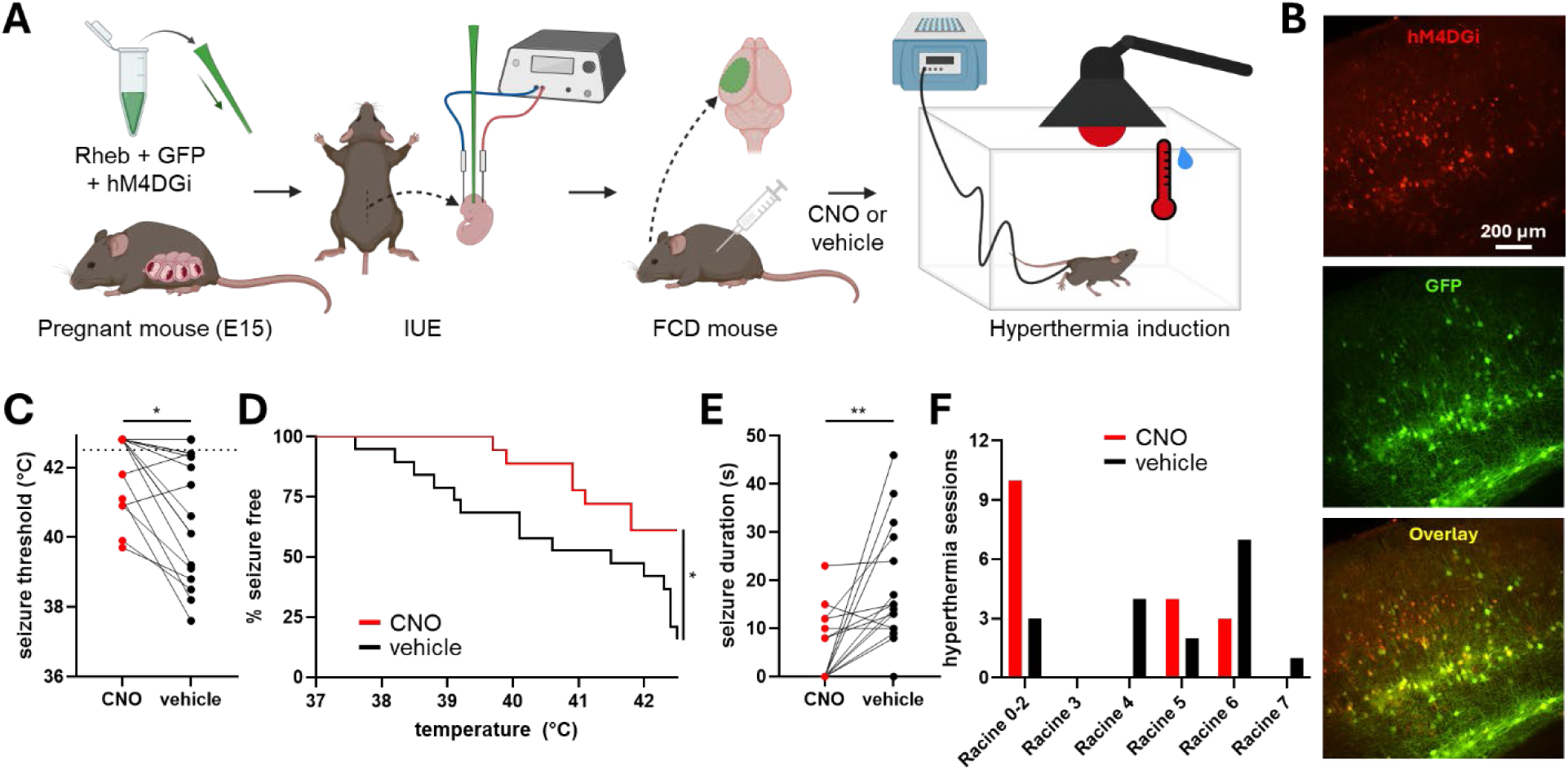
Chemogenetic inhibition of excitatory neurons ameliorates hyperthermia-evoked seizures in *Rheb-*FCD mice. **(A)** Cartoon illustrating experimental overview for chemogenetic manipulation of hyperthermia-evoked seizures. Rheb, GFP, and the inhibitory DREADD, hM4DGi, are introduced via IUE. Mice are exposed to hyperthermia following an injection with either CNO (to activate hM4DGi) or vehicle control. **(B)** Expression of hM4DGi (red) within the dysplastic region (as delineated by GFP; green). **(C)** Mice treated with CNO exhibit a higher seizure temperature threshold. Note that in this graph, data points above the experimental end-point (42.5 °C) indicate a trial in which no seizure was elicited. Seizures were elicited in only 39% of trials with CNO, versus 84% of trials with vehicle (p = 0.015, Fisher’s exact test, n = 18-19 mice). In trials in which seizures did occur, mean threshold was 40.9 ± 0.3 °C for CNO (n = 7) versus 40.3 ± 0.5 °C for vehicle (n = 14). **(D)** Seizure thresholds across all sessions were plotted as a Kaplan-Meier curve to capture both threshold data and the percentage of mice that exhibited seizures. CNO reduced seizures (p = 0.0173, log-rank (Mantel-Cox) test). **(E)** Seizure duration with or without CNO. When sessions in which no seizures occurred were coded as duration = 0, CNO significantly reduced seizure duration (p = 0.0015, Wilcoxon match-pairs signed rank test). Excluding sessions for which no seizures occurred, mean seizure duration was 13 ± 2 seconds for CNO (n = 7) versus 20 ± 3 seconds for vehicle (n = 14). **(F)** Distribution of maximum Racine scores with or without CNO.

Using co-electroporation (together with pCAG-RhebP37L and pCAG-GFP), we introduced hM4DGi, an inhibitory Designer Receptor Exclusively Activated by Designer Drugs (DREADD) (Armbruster et al., 2007), under control of the excitatory neuron promoter, CaMKIIα (Liu and Jones, 1996). We confirmed co-expression of hM4DGi and GFP at the regional level: i.e., hM4DGi was expressed within the dysplastic region (Fig. 4B). To further confirm that hM4DGi was expressed within neurons that were activated by hyperthermia-evoked seizures, we evoked seizures (in the absence of CNO) and quantified the co-localization between c-Fos and hM4DGi. We found hM4DGi expression within approximately ∼40% of c-Fos+ cells (quantified across a total of n = 571 c-Fos+ cells / 6 mice). Next, to test the impact of chemogenetic inhibition on seizures, we exposed the animals to hyperthermia and recorded the seizure temperature threshold, duration, and severity. We found that pre-administration of the ligand Clozapine-N-Oxide (CNO) was significantly protective against seizures evoked by hyperthermia, as demonstrated by fewer / an increased threshold for hyperthermia-evoked seizures (Fig. 4C-F). As a negative control, we confirmed that CNO had no direct impact on seizure threshold on *Rheb-*FCD mice that did not express hM4DGi (Supplementary Fig. 3).

For hyperthermia to serve as an effective screening platform in the *Rheb-*FCD model, the same manipulations that ameliorate acute evoked seizures should similarly ameliorate spontaneous ictal activity and interictal abnormalities. We tested this by conducting continuous EEG recordings across 6 days, delivering once daily alternating (cross-balanced) injections of CNO versus vehicle (Fig. 5A). To identify interictal spikes, we optimized an automated spike detector – with detections based upon amplitude, width, prominence, and interspike interval – which we validated against manual human spike counts across 20 randomly selected 15-minute bins. To identify spontaneous seizures, we manually reviewed the EEG recordings (blinded to drug condition) to identify electrographic seizures. We then used a custom-coded automated detector of sustained periods of high-amplitude activity to guide a second, targeted review of the EEG data. In total, we identified 215 seizures across ∼1584 aggregate “mouse hours” of recording: the first round of manual review identified 211 of these (i.e., 98.1% sensitivity), with the additional 4 seizures identified on the second round of assisted review. Our detector, which had been deliberately tuned for high sensitivity, correctly identified 203/215 seizures (i.e., 94.4% sensitivity), with a false alarm rate of 0.41 false detections per mouse hour of recording.

**Figure 5.**
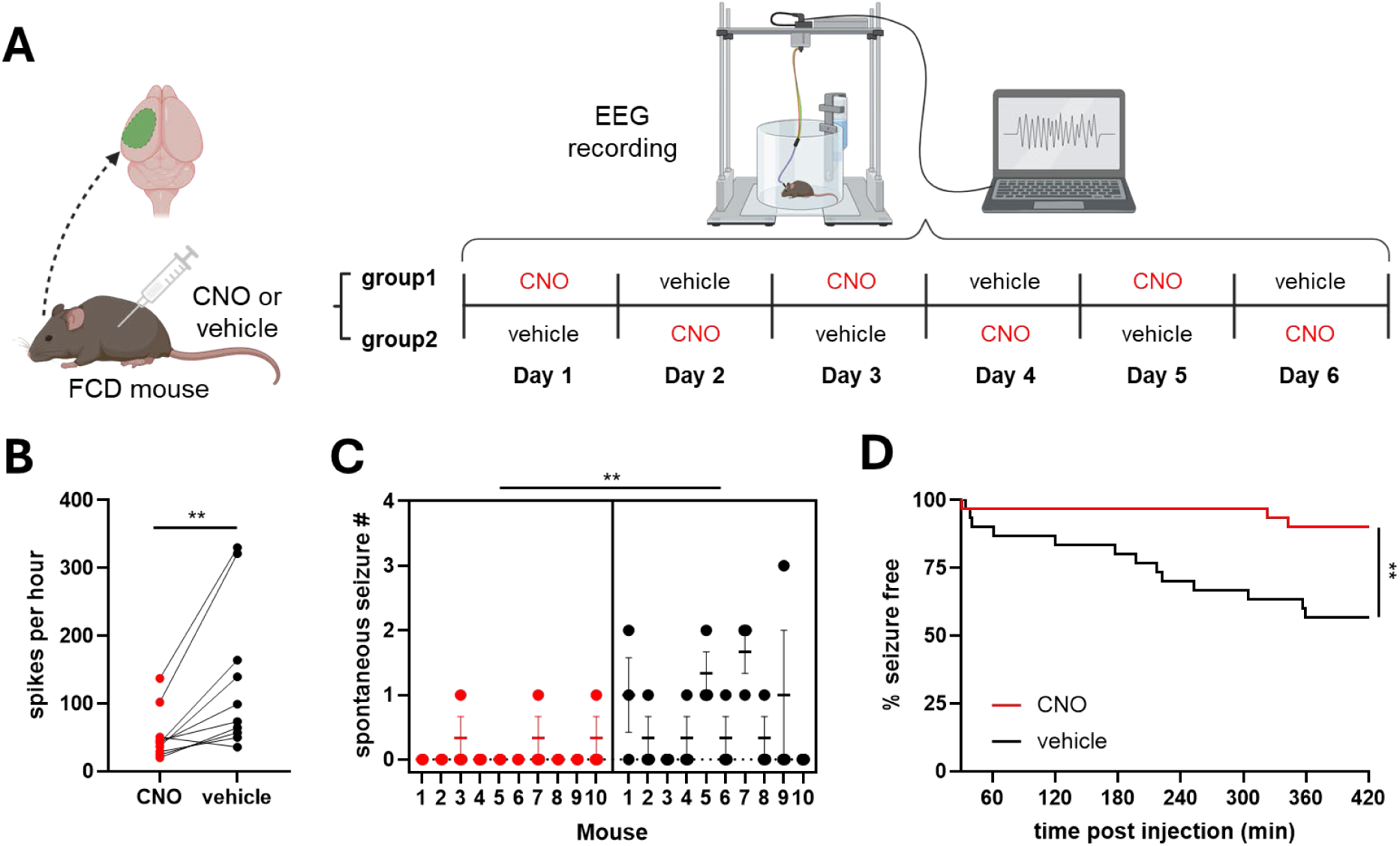
Chemogenetic inhibition of excitatory neurons ameliorates spontaneous seizures in *Rheb-*FCD mice. **(A)** Cartoon illustrating experimental overview for chemogenetic manipulation of spontaneous seizures. Chronic EEG recordings are performed for 6 sequential days. Mice are assigned counter-balanced groups injected once daily with either CNO or a vehicle control. **(B)** Interictal spike counts were averaged within and then across mice for each condition. CNO significantly reduced the frequency of interictal spikes (53 ± 12 per hour with CNO versus 134 ± 34 per hour with vehicle; p = 0.009, paired t-test). n = 10 mice. **(C)** Number of spontaneous seizures occurring within a time window of 30-360 minutes post injection. CNO significantly reduced the occurrence of seizures (p = 0.01, two-tailed nested t-test). n = 10 mice; 6 sessions per mouse (3 CNO / 3 vehicle). **(D)** Seizure occurrence was plotted as a Kaplan-Meier curve as a function of time post injection with either CNO or vehicle. CNO significantly reduced the occurrence of seizures (p = 0.004, log-rank (Mantel-Cox) test).

We narrowed our analysis window to 30-240 minutes post-injection (during which there should be ongoing DREADD activation (Alexander et al., 2009; Krashes et al., 2011)) and quantified interictal spikes and seizures within that window. We observed a significant reduction of both interictal spike burden and spontaneous seizure occurrence in the CNO-treated condition relative to vehicle control (Fig. 5B-D).

In summary, our results suggest that chemogenetic inhibition of excitatory neurons within the dysplasia ameliorates hyperthermia-evoked seizures, as well as spontaneous interictal spikes and seizures in *Rheb-*FCD mice.

## Discussion

The ability to acutely evoke seizures in experimental animal models – and to do so in a manner with high construct, face, and predictive validity – will facilitate advances in our mechanistic understanding of seizures and our development of new therapeutic tools in epilepsy. In this study, we established that acute elevation of body temperature (i.e., hyperthermia) could serve as a reliable seizure trigger in a *Rheb*-FCD mouse model. We then used this induced seizure paradigm to demonstrate that chemogenetic inhibition of excitatory neurons within the dysplastic region had a significant anti-seizure effect. Finally, we confirmed the predictive validity of this acute result by applying the same manipulation chronically to demonstrate similar inhibition of interictal spikes and spontaneous seizures. Together, these results establish a new, acute paradigm for induction of acute seizures in an animal model of a common refractory focal epilepsy.

Our central finding is that hyperthermia reliably and reproducibly provokes electrographic and behavioral seizures in adult *Rheb*-FCD mice (Figs. 1-3, Supplementary Fig. 2). In fact, the mean seizure threshold in the *Rheb*-FCD mice was comparable to that observed in *Scn1a^+/−^*mice (Fig. 1B-C), in which the hyperthermia-evoked seizure phenotype is already well established (Oakley et al., 2009; Mistry et al., 2014). To our knowledge, this is the first demonstration that hyperthermia can trigger seizures in a focal, non-genetic epilepsy model. The generalizability of this phenotype could be tested across other common models of focal epilepsy – such as temporal lobe epilepsy (Lévesque et al., 2016) – using a similar experimental design.

Why does hyperthermia evoke seizures in the *Rheb*-FCD mice? Indeed, the mechanism by which fever triggers breakthrough seizures in persons with epilepsy (Bauer et al., 2000; Balamurugan et al., 2013; Kaddumukasa et al., 2013; Illingworth et al., 2014; Wassenaar et al., 2014; Kurian et al., 2018; Rafati et al., 2023), including FCD (Singh et al., 2001; Itamura et al., 2019), remains incompletely understood. Brain temperature directly alters neuronal excitability and firing (e.g., (Shibasaki et al., 2007; Kim and Connors, 2012; Burek et al., 2019), which may interact with underlying pathology in *Rheb*-FCD mice to increase the probability of epileptiform activity. In the clinical context, hyperthermia typically occurs alongside a broader inflammatory response, which independently impacts neurotransmission (Mosili et al., 2020). Finally, we anecdotally noted that the *Rheb*-FCD mice tended to have seizures when being actively handled, suggesting that they may be broadly susceptible to stress. Future studies will be required to systematically test whether other paradigms, such as restraint stress (Molina et al., 2023), can reliably evoke seizures, but regardless, hyperthermia is experimentally advantageous in that it is easily titratable, quantifiable, and reproducible trial-to-trial.

We additionally observed that hyperthermia-evoked seizures result in significant c-Fos expression within the FCD (Fig. 3). Notably, our experimental design did not provide cellular resolution of RHEB+ versus RHEB-cells within the dysplastic region – as RHEB and GFP were delivered via separate plasmids and thus were likely introduced into imperfectly overlapping neuronal populations – so we have not attempted to quantify the relative c-Fos expression in RHEB+ versus RHEB-neurons. Future experiments could achieve more selective labeling by combining (1) a single plasmid to introduce both Rheb and Cre, with (2) a Cre-dependent GFP. Fluorescent calcium indicators (e.g., GCaMP (Dana et al., 2019)) could alternatively be introduced (within any desired neuronal population, within or beyond the FCD) to measure neuronal activity at much higher temporal resolution. Indeed, the acute nature of hyperthermia-evoked seizures should facilitate fiber photometry recording or live imaging timed to the peri-ictal period (Somarowthu et al., 2021).

Finally, we found that chemogenetic inhibition of a subset of excitatory neurons within the FCD significantly protects against hyperthermia-evoked seizures (Fig. 4) and confirmed a similar protection against interictal spikes and spontaneous seizures in chronic recordings (Fig. 5). These findings are in line with recent targeted approaches that similarly improve the epilepsy phenotype in FCD models, including inhibition of a hyperpolarization-activated cyclic nucleotide–gated potassium channel (HCN4) ectopically-expressed within dysplastic cells (Hsieh et al., 2020), and viral over-expression of K_v_1.1 potassium channels in CaMKIIα-expressing cells within the FCD (Almacellas Barbanoj et al., 2024). Thus, our work provides further pre-clinical justification for development of spatially-targeted molecular therapies for treatment of FCD.

More broadly, by establishing acute hyperthermia as a reliable and modifiable seizure trigger in the well-established *Rheb*-FCD model, we have identified a screening paradigm that is significantly higher-throughput than the chronic recordings required to detect changes in spontaneous seizure rate. Furthermore, since hyperthermia can be administered repeatedly, therapies can be screened using a within-subject design across consecutive trials, enhancing statistical power and minimizing animal use. Thus, this acute model could be used to accelerate the preclinical developmental pipeline for new antiseizure therapies for refractory focal epilepsy (e.g., the NINDS Epilepsy Therapy Screening Program (ETSP) (Kehne et al., 2017)).

## CRedIT authorship contribution statement

Joanna Mattis and Yu Wang contributed to project conceptualization. Joanna Mattis, Xiaogang Zhang, Ruitao Shen, and Chandni Rana contributed to data curation. All authors contributed to formal data analysis. Joanna Mattis contributed to funding acquisition. Xiaogang Zhang, Chandni Rana, Ruitao Shen, Tao Yang, Joseph Barden, Hudson Haberland, Mi Jiang, and Rida Qureshi contributed to the research investigation process, that is, performing experiments and collecting data. Joanna Mattis was responsible for project administration and management. Saif Siddiqui, Alexander Sajdak, and Patrick Lawlor contributed to programming, software development, and implementation of analysis code. Joseph Barden, Yu Wang, and Joanna Mattis contributed to the provision of resources including study materials, reagents, and animals. Joanna Mattis was responsible for oversight and leadership supervision. Xiaogang Zhang, Chandni Rana, Ruitao Shen, Joseph Barden, Mi Jiang, Yu Wang, and Joanna Mattis contributed to data visualization. Xiaogang Zhang, Ruitao Shen, and Joanna Mattis wrote the original draft of the manuscript, with contributions from Chandni Rana, Joseph Barden, Hudson Haberland, Saif Siddiqui, and Alexander Sajdak. All authors contributed to reviewing and editing the manuscript.

## Ethical statement

All animal procedures were approved by the Institutional Animal Care and Use Committee at the University of Michigan and were conducted in accordance with the United States Public Health Service’s Policy on Humane Care and Use of Laboratory Animals.

## Acknowledgments

Research reported in this publication was supported by the National Institute of Neurological Disorders and Stroke (NINDS) of the National Institutes of Health (NIH), under Award Number T32 NS115724 (C.R.), 1F31 NS14319501 (C.R.), K08 NS121464 (J.M), 5K12 HD028820-33 (P.L.), and R01NS136181 (Y.W.). Additional support was obtained from a Kenneth Eisenberg Emerging Scholar award from the Taubman Institute at the University of Michigan (J.M.). We thank all members of the Mattis lab for helpful discussion. The content is solely the responsibility of the authors and does not necessarily represent the official views of the NIH.

## References

1. Alexander GM, Rogan SC, Abbas AI, Armbruster BN, Pei Y, Allen JA, Nonneman RJ, Hartmann J, Moy SS, Nicolelis MA, McNamara JO, Roth BL (2009) Remote control of neuronal activity in transgenic mice expressing evolved G protein-coupled receptors. Neuron 63:27–39.

2. Almacellas Barbanoj A, Graham RT, Maffei B, Carpenter JC, Leite M, Hoke J, Hardjo F, Scott-Solache J, Chimonides C, Schorge S, Kullmann DM, Magloire V, Lignani G (2024) Anti-seizure gene therapy for focal cortical dysplasia. Brain 147:542–553.

3. Armbruster BN, Li X, Pausch MH, Herlitze S, Roth BL (2007) Evolving the lock to fit the key to create a family of G protein-coupled receptors potently activated by an inert ligand. Proc Natl Acad Sci U S A 104:5163–5168.

4. Balamurugan E, Aggarwal M, Lamba A, Dang N, Tripathi M (2013) Perceived trigger factors of seizures in persons with epilepsy. Seizure 22:743–747.

5. Bauer J, Saher MS, Burr W, Elger CE (2000) Precipitating factors and therapeutic outcome in epilepsy with generalized tonic-clonic seizures. Acta Neurol Scand 102:205–208.

6. Blumcke I et al. (2017) Histopathological Findings in Brain Tissue Obtained during Epilepsy Surgery. N Engl J Med 377:1648–1656.

7. Burek M, Follmann R, Rosa E (2019) Temperature effects on neuronal firing rates and tonic-to-bursting transitions. Biosystems 180:1–6.

8. Cavalheiro EA (1995) The pilocarpine model of epilepsy. Ital J Neurol Sci 16:33–37.

9. Chen Z, Brodie MJ, Liew D, Kwan P (2018) Treatment Outcomes in Patients With Newly Diagnosed Epilepsy Treated With Established and New Antiepileptic Drugs: A 30-Year Longitudinal Cohort Study. JAMA Neurol 75:279–286.

10. Chern JJ, Patel AJ, Jea A, Curry DJ, Comair YG (2010) Surgical outcome for focal cortical dysplasia: an analysis of recent surgical series. J Neurosurg Pediatr 6:452–458.

11. Cohen-Gadol AA, Özduman K, Bronen RA, Kim JH, Spencer DD (2004) Long-term outcome after epilepsy surgery for focal cortical dysplasia. Journal of Neurosurgery 101:55–65.

12. Connolly MB (2016) Dravet Syndrome: Diagnosis and Long-Term Course. Can J Neurol Sci 43 Suppl 3:S3–8.

13. Crino PB (2009) Focal brain malformations: seizures, signaling, sequencing. Epilepsia 50 Suppl 9:3–8.

14. Cwetsch AW, Ziogas I, Narducci R, Savardi A, Bolla M, Pinto B, Perlini LE, Bassani S, Passafaro M, Cancedda L (2022) A rat model of a focal mosaic expression of PCDH19 replicates human brain developmental abnormalities and behaviours. Brain Commun 4:fcac091.

15. Dana H, Sun Y, Mohar B, Hulse BK, Kerlin AM, Hasseman JP, Tsegaye G, Tsang A, Wong A, Patel R, Macklin JJ, Chen Y, Konnerth A, Jayaraman V, Looger LL, Schreiter ER, Svoboda K, Kim DS (2019) High-performance calcium sensors for imaging activity in neuronal populations and microcompartments. Nat Methods 16:649–657.

16. Dravet C, Oguni H (2013) Dravet syndrome (severe myoclonic epilepsy in infancy). Handb Clin Neurol 111:627–633.

17. Erum JV, Dam DV, Deyn PPD (2019) PTZ-induced seizures in mice require a revised Racine scale. Epilepsy Behav 95:51–55.

18. Fiest KM, Sauro KM, Wiebe S, Patten SB, Kwon C-S, Dykeman J, Pringsheim T, Lorenzetti DL, Jetté N (2017) Prevalence and incidence of epilepsy: A systematic review and meta-analysis of international studies. Neurology 88:296–303.

19. Giardina WJ, Gasior M (2009) Acute seizure tests in epilepsy research: electroshock– and chemical-induced convulsions in the mouse. Curr Protoc Pharmacol Chapter 5:Unit 5.22.

20. Griffin A, Hamling KR, Hong S, Anvar M, Lee LP, Baraban SC (2018) Preclinical Animal Models for Dravet Syndrome: Seizure Phenotypes, Comorbidities and Drug Screening. Front Pharmacol 9:573.

21. Gu B, Zhu M, Glass MR, Rougié M, Nikolova VD, Moy SS, Carney PR, Philpot BD (2019) Cannabidiol attenuates seizures and EEG abnormalities in Angelman syndrome model mice. J Clin Invest 129:5462–5467.

22. Guerrini R, Dobyns WB (2014) Malformations of cortical development: clinical features and genetic causes. Lancet Neurol 13:710–726.

23. Harvey AS, Cross JH, Shinnar S, Mathern GW, ILAE Pediatric Epilepsy Surgery Survey Taskforce (2008) Defining the spectrum of international practice in pediatric epilepsy surgery patients. Epilepsia 49:146–155.

24. Hsieh LS, Wen JH, Claycomb K, Huang Y, Harrsch FA, Naegele JR, Hyder F, Buchanan GF, Bordey A (2016) Convulsive seizures from experimental focal cortical dysplasia occur independently of cell misplacement. Nat Commun 7:11753.

25. Hsieh LS, Wen JH, Nguyen LH, Zhang L, Getz SA, Torres-Reveron J, Wang Y, Spencer DD, Bordey A (2020) Ectopic HCN4 expression drives mTOR-dependent epilepsy in mice. Sci Transl Med 12:eabc1492.

26. Hudson AE (2018) Genetic reporters of neuronal activity: c-Fos and G-CaMP6. Methods Enzymol 603:197–220.

27. Illingworth JL, Watson P, Ring H (2014) Why do seizures occur when they do? Situations perceived to be associated with increased or decreased seizure likelihood in people with epilepsy and intellectual disability. Epilepsy Behav 39:78–84.

28. Itamura S, Okanishi T, Arai Y, Nishimura M, Baba S, Ichikawa N, Hirayama Y, Ishihara N, Hiraide T, Ishigaki H, Fukuda T, Otsuki Y, Enoki H, Fujimoto A (2019) Three Cases of Hemiconvulsion-Hemiplegia-Epilepsy Syndrome With Focal Cortical Dysplasia Type IIId. Front Neurol 10 Available at: https://www.frontiersin.org/journals/neurology/articles/10.3389/fneur.2019.01233/full [Accessed November 11, 2025].

29. Jafarian M, Esmaeil Alipour M, Karimzadeh F (2020) Experimental Models of Absence Epilepsy. Basic Clin Neurosci 11:715–726.

30. Jayalakshmi S, Nanda SK, Vooturi S, Vadapalli R, Sudhakar P, Madigubba S, Panigrahi M (2019) Focal Cortical Dysplasia and Refractory Epilepsy: Role of Multimodality Imaging and Outcome of Surgery. American Journal of Neuroradiology 40:892–898.

31. Kaddumukasa M, Kaddumukasa M, Matovu S, Katabira E (2013) The frequency and precipitating factors for breakthrough seizures among patients with epilepsy in Uganda. BMC Neurol 13:182.

32. Kao H-Y, Yao Y, Yang T, Ziobro J, Zylinski M, Mir MY, Hu S, Cao R, Borna NN, Banerjee R, Parent JM, Wang S, Leventhal DK, Li P, Wang Y (2023) Sudden Unexpected Death in Epilepsy and Respiratory Defects in a Mouse Model of DEPDC5-Related Epilepsy. Ann Neurol 94:812–824.

33. Karler R, Murphy V, Calder LD, Turkanis SA (1989) Pentylenetetrazol kindling in mice. Neuropharmacology 28:775–780.

34. Kehne JH, Klein BD, Raeissi S, Sharma S (2017) The National Institute of Neurological Disorders and Stroke (NINDS) Epilepsy Therapy Screening Program (ETSP). Neurochem Res 42:1894–1903.

35. Kim J, Connors B (2012) High temperatures alter physiological properties of pyramidal cells and inhibitory interneurons in hippocampus. Front Cell Neurosci 6 Available at: https://www.frontiersin.org/journals/cellular-neuroscience/articles/10.3389/fncel.2012.00027/full [Accessed December 15, 2025].

36. Krashes MJ, Koda S, Ye C, Rogan SC, Adams AC, Cusher DS, Maratos-Flier E, Roth BL, Lowell BB (2011) Rapid, reversible activation of AgRP neurons drives feeding behavior in mice. J Clin Invest 121:1424–1428.

37. Kravchenko JA, Goldberg EM, Mattis J (2023) Optogenetic and chemogenetic manipulation of seizure threshold in mice. STAR Protoc 4:102019.

38. Kurian M, Korff CM, Ranza E, Bernasconi A, Lübbig A, Nangia S, Ramelli GP, Wohlrab G, Nordli DR, Bast T (2018) Focal cortical malformations in children with early infantile epilepsy and PCDH19 mutations: case report. Dev Med Child Neurol 60:100–105.

39. Kwan P, Arzimanoglou A, Berg AT, Brodie MJ, Allen Hauser W, Mathern G, Moshé SL, Perucca E, Wiebe S, French J (2010) Definition of drug resistant epilepsy: consensus proposal by the ad hoc Task Force of the ILAE Commission on Therapeutic Strategies. Epilepsia 51:1069–1077.

40. Leech CK, McIntyre DC (1976) Kindling rates in inbred mice: an analog to learning? Behavioral Biology 16:439–452.

41. Lévesque M, Avoli M (2013) The kainic acid model of temporal lobe epilepsy. Neurosci Biobehav Rev 37:2887–2899.

42. Lévesque M, Avoli M, Bernard C (2016) Animal models of temporal lobe epilepsy following systemic chemoconvulsant administration. J Neurosci Methods 260:45–52.

43. Liu XB, Jones EG (1996) Localization of alpha type II calcium calmodulin-dependent protein kinase at glutamatergic but not gamma-aminobutyric acid (GABAergic) synapses in thalamus and cerebral cortex. Proceedings of the National Academy of Sciences 93:7332–7336.

44. Marshall GF, Gonzalez-Sulser A, Abbott CM (2021) Modelling epilepsy in the mouse: challenges and solutions. Dis Model Mech 14:dmm047449.

45. Mattis J, Somarowthu A, Goff KM, Jiang E, Yom J, Sotuyo N, Mcgarry LM, Feng H, Kaneko K, Goldberg EM (2022) Corticohippocampal circuit dysfunction in a mouse model of Dravet syndrome. Elife 11:e69293.

46. Meisler MH (2019) SCN8A encephalopathy: Mechanisms and models. Epilepsia 60 Suppl 3:S86–S91.

47. Mistry AM, Thompson CH, Miller AR, Vanoye CG, George AL, Kearney JA (2014) Strain– and age-dependent hippocampal neuron sodium currents correlate with epilepsy severity in Dravet syndrome mice. Neurobiol Dis 65:1–11.

48. Molina P, Andero R, Armario A (2023) Restraint or immobilization: A comparison of methodologies for restricting free movement in rodents and their potential impact on physiology and behavior. Neuroscience & Biobehavioral Reviews 151:105224.

49. Mosili P, Maikoo S, Mabandla M, Vuyisile, Qulu L (2020) The Pathogenesis of Fever-Induced Febrile Seizures and Its Current State. Neurosci Insights 15:2633105520956973.

50. Nguyen LH, Bordey A (2022) Current Review in Basic Science: Animal Models of Focal Cortical Dysplasia and Epilepsy. Epilepsy Curr 22:234–240.

51. Nguyen LH, Mahadeo T, Bordey A (2019) mTOR Hyperactivity Levels Influence the Severity of Epilepsy and Associated Neuropathology in an Experimental Model of Tuberous Sclerosis Complex and Focal Cortical Dysplasia. J Neurosci 39:2762–2773.

52. Oakley JC, Kalume F, Yu FH, Scheuer T, Catterall WA (2009) Temperature– and age-dependent seizures in a mouse model of severe myoclonic epilepsy in infancy. Proc Natl Acad Sci U S A 106:3994–3999.

53. Proietti Onori M, Koene LMC, Schäfer CB, Nellist M, de Brito van Velze M, Gao Z, Elgersma Y, van Woerden GM (2021) RHEB/mTOR hyperactivity causes cortical malformations and epileptic seizures through increased axonal connectivity. PLoS Biol 19:e3001279.

54. Rafati A, Jameie M, Amanollahi M, Jameie M, Pasebani Y, Sakhaei D, Ilkhani S, Rashedi S, Pasebani MY, Azadi M, Rahimlou M, Kwon C-S (2023) Association of seizure with COVID-19 vaccines in persons with epilepsy: A systematic review and meta-analysis. J Med Virol 95:e29118.

55. Ramos-Lizana J, Rodriguez-Lucenilla MI, Aguilera-López P, Aguirre-Rodríguez J, Cassinello-García E (2012) A study of drug-resistant childhood epilepsy testing the new ILAE criteria. Seizure 21:266–272.

56. Schmidt J (1987) Changes in seizure susceptibility in rats following chronic administration of pentylenetetrazol. Biomed Biochim Acta 46:267–270.

57. Scott KEJ, Hermosillo Arrieta MF, Williams AJ (2025) Deciphering SCN2A: A comprehensive review of rodent models of Scn2a dysfunction. Epilepsia.

58. Shibasaki K, Suzuki M, Mizuno A, Tominaga M (2007) Effects of Body Temperature on Neural Activity in the Hippocampus: Regulation of Resting Membrane Potentials by Transient Receptor Potential Vanilloid 4. J Neurosci 27:1566–1575.

59. Singh G, Dhanuka AK, Bains HS, Singh D (2001) Febrile Status Epilepticus as the First Presentation of Cortical Developmental Malformation: Report of 2 Cases. Neurology India 49:287.

60. Somarowthu A, Goff KM, Goldberg EM (2021) Two-photon calcium imaging of seizures in awake, head-fixed mice. Cell Calcium 96:102380.

61. Wassenaar M, Kasteleijn-Nolst Trenité DGA, de Haan G-J, Carpay JA, Leijten FSS (2014) Seizure precipitants in a community-based epilepsy cohort. J Neurol 261:717–724.

62. Yan L, Findlay GM, Jones R, Procter J, Cao Y, Lamb RF (2006) Hyperactivation of mammalian target of rapamycin (mTOR) signaling by a gain-of-function mutant of the Rheb GTPase. J Biol Chem 281:19793–19797.

