## Supplemental figures for "Characterization and application of hyperthermia-evoked seizures in a mouse model of focal cortical dysplasia"

### Supplementary figures

**Supplementary Figure 1. IUE-mediated somatic mutagenesis creates mice with an FCD phenotype.**

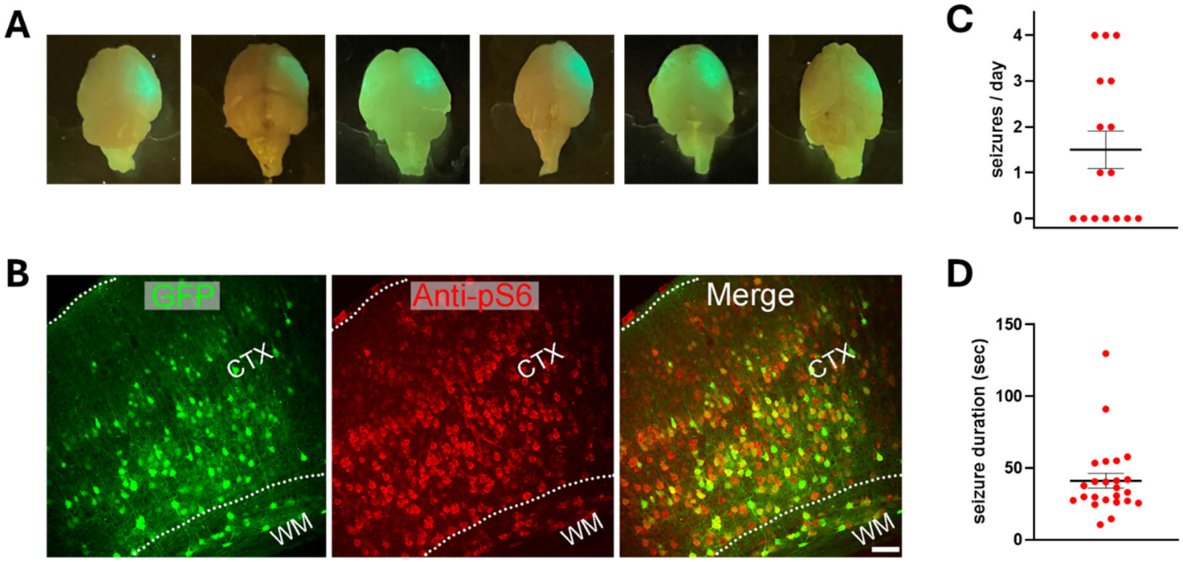


**(A)** Gross visualization of electroporated region as seen under a fluorescent flashlight.

**(B)** *Rheb* IUE induces mTOR hyperactivation (pS6, red) in transfected neurons (GFP). CTX: cortex; WM: white matter. Scale bar represents 100 μm.

**(C)** FCD mice exhibited spontaneous electrographic seizures (mean 1.5 ± 0.4 seizures per day; n = 16 days across 10 mice).

**(D)** Electrographic seizure durations ranged from ~20-130 seconds (mean 41 ± 5 seconds; n = 24 seizures across 10 mice).

**Supplementary Figure 2. Hyperthermia-evoked seizures in *Rheb*-FCD mice are robust across sex and strain.**

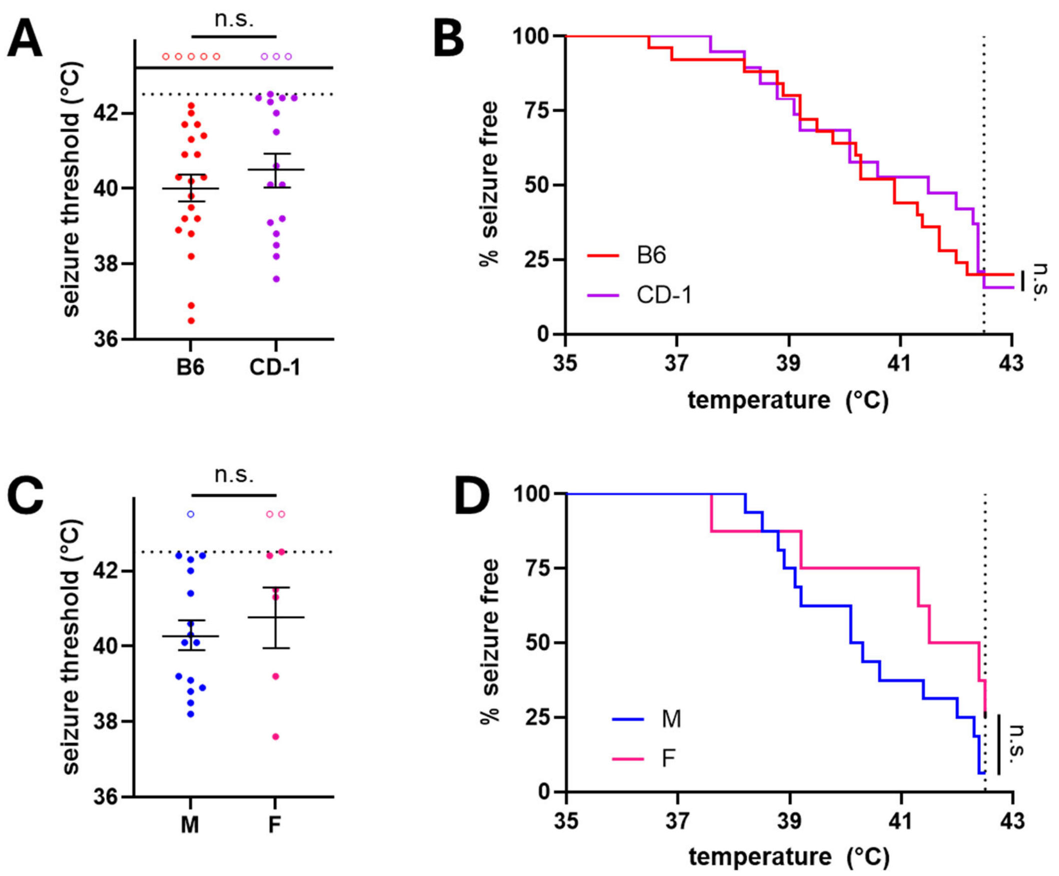


**(A)** Hyperthermia-evoked seizure thresholds for *Rheb*-FCD mice were similar between mice on a C57BL/6 background (red circles) and those on a CD-1 background (purple circles). For sessions that did elicit seizures, mean seizure threshold was similar between groups (40.0 ± 0.4 °C for C57BL/6 mice versus 40.5 ± 0.4 °C for CD-1 mice, p = 0.39, unpaired t-test). n = 5 mice / 20 seizures / 25 sessions for C57BL/6 and 19 mice / 16 seizures / 19 sessions for CD-1.

**(B)** Seizure thresholds across all sessions were potted as a Kaplan-Meier curve to capture both threshold data and the percentage of mice that exhibited seizures. C57BL/6 versus CD-1 curves were similar (p = 0.76, log-rank (Mantel-Cox) test).

**(C)** Hyperthermia-evoked seizure thresholds for *Rheb*-FCD mice were similar between male mice (blue circles) and female mice (pink circles). All thresholds were from first seizure sessions pooled across strains. For sessions that did elicit seizures, mean seizure threshold was 40.3 ± 0.4 °C for male mice versus 40.8 ± 0.8 °C for female mice, p = 0.56, unpaired t-test). n = 16 mice / 15 seizures for male mice and 8 mice / 6 seizures for female mice.

**(D)** Seizure thresholds across all sessions were plotted as a Kaplan-Meier curve to capture both threshold data and the percentage of mice that exhibited seizures. Male versus female curves were similar (p = 0.13, log-rank (Mantel-Cox) test).

**Supplementary Figure 3. CNO administration alone does not impact hyperthermia-evoked seizure threshold.**

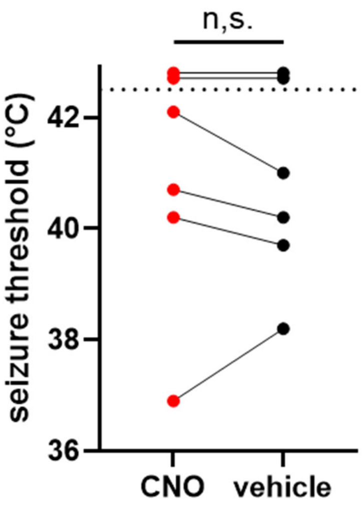


Seizure threshold of *Rheb*-FCD mice that did not express hM4DGi, with versus without pre-treatment with CNO (n = 6). Note that in this graph, data points above the experimental end-point (42.5 °C) indicate a trial in which no seizure was elicited. In trials in which seizures did occur, mean threshold was 41.1 ± 0.7 °C for CNO versus 40.8 ± 0.4 °C for vehicle (p = 0.39, paired t-test).
